# Distinct roles of glutamine metabolism in benign and malignant cartilage tumors with *IDH* mutations

**DOI:** 10.1101/2021.09.12.459996

**Authors:** Hongyuan Zhang, Vijitha Puviindran, Puviindran Nadesan, Xiruo Ding, Leyao Shen, Yuning J. Tang, Hidetoshi Tsushima, Yasuhito Yahara, Ga I Ban, Guo-Fang Zhang, Courtney M. Karner, Benjamin Alman

## Abstract

Enchondromas and chondrosarcomas are common cartilage neoplasms that are either benign or malignant respectively. The majority of these tumors harbor mutations in either *IDH1* or *IDH2.* Glutamine metabolism has been implicated as a critical regulator of tumors with *IDH* mutations. Chondrocytes and chondrosarcomas with mutations in the *IDH1* or *IDH2* genes showed enhanced glutamine utilization in downstream metabolism. Using genetic and pharmacological approaches, we demonstrated that glutaminase-mediated glutamine metabolism played distinct roles in enchondromas and chondrosarcomas with *IDH1* or *IDH2* mutations. Deletion of glutaminase in chondrocytes with *Idh1* mutation increased the number and size of enchondroma-like lesions. Pharmacological inhibition of glutaminase in chondrosarcoma xenografts reduced overall tumor burden. Glutamine affected cell differentiation and viability in these tumors differently through different downstream metabolites. During murine enchondroma-like lesion development, glutamine-derived *α*-ketoglutarate promoted hypertrophic chondrocyte differentiation and regulated chondrocyte proliferation. In human chondrosarcoma, glutamine-derived non-essential amino acids played an important role in preventing cell apoptosis. This study reveals that glutamine metabolism can play distinct roles in benign and malignant cartilage tumors sharing the same genetic mutations. Inhibiting GLS may provide a therapeutic approach to suppress chondrosarcoma tumor growth.

## Introduction

Enchondroma is a common benign cartilaginous neoplasm and is estimated to be present in 3% of the total population [1, 2]. These tumors develop from dysregulated chondrocyte differentiation in the growth plate and are mostly present in the metaphysis of long bones [3]. In patients with multiple enchondromatosis (more than one enchondroma lesions) such as Maffucci׳s syndrome and Ollier׳s disease, the risk of malignant transformation is reported to be up to 60% [4]. Chondrosarcoma is the second most common primary malignancy of the bone [4]. They arise *de novo* or develop from preexisting benign tumors including enchondromas [4]. High-grade chondrosarcomas have high metastatic potential and poor prognosis [5]. Currently there are no universally effective pharmacologic therapies for enchondromas or chondrosarcomas.

Somatic mutations of isocitrate dehydrogenase 1 and 2 (*IDH1* and *IDH2*) are the most frequent genetic variations in enchondromas and chondrosarcomas [6–10]. They are present in 56% - 90% of enchondroma tumors [6, 8], and in about 50% of chondrosarcoma tumors [7, 9]. Although *IDH1* or *IDH2* mutations are present in enchondromas and chondrosarcomas, it is not known whether these tumors share similar metabolic requirements for tumor development and cell viability. Wildtype IDH1 and IDH2 enzymes catalyze the reversible conversion between isocitrate and *α*- ketoglutarate (*α*-KG) in the cytoplasm or the mitochondria respectively. Mutant IDH 1 or IDH2 enzymes lose their original function and gain a neomorphic function that converts *α*-KG to D-2-hydroxyglutarate (D-2HG) [11–13]. D-2HG is considered as a putative “oncometabolite” in various cancers with mutations in *IDH1/2* [14–20]. Interestingly, pharmacological inhibition of mutant IDH1 enzyme did not alter chondrosarcoma tumorigenesis despite effective reduction of D-2HG synthesis [21]. Several clinical trials of mutant IDH inhibitors have been conducted in patients with chondrosarcoma [22].

The results to date have been variable, showing at best stabilization of the disease [23]. Recently, an investigation of the metabolomes in chondrosarcomas showed global alterations in cellular metabolism in *IDH1* or *IDH2* mutant chondrosarcomas, suggesting such alteration might drive the neoplastic phenotype, or be possible therapeutic targets for cancers with *IDH1* or *IDH2* mutations [24].

Glutamine metabolism is an important metabolic pathway that is critical for the survival of various cancers as well as proper proliferation and differentiation of different cell types [25–31]. Glutamine metabolism starts when glutaminase (GLS) deaminates glutamine to glutamate. Glutamate could be further used to generate *α*-ketoglutarate (*α*- KG), glutathione, other non-essential amino acids, and nucleotides, etc. Through different downstream metabolites, glutamine regulates cancer cell behaviors by modulating bioenergetics, biosynthesis, redox homeostasis, etc. [26]. In cancers with *IDH1* or *IDH2* mutations, glutamine is utilized as the primary source for D-2HG production [11, 32, 33]. In addition, some tumors with *IDH1* or *IDH2* mutations are reported to be dependent on glutamine metabolism for tumor growth or cell viability [34–37]. In non-cancerous cells, glutamine metabolism regulates cell differentiation mainly through many downstream metabolites including amino acids as well as *α*-KG and acetyl-CoA which act as cofactors for various histone modifying enzymes [28, 30, 38, 39].

In the context of cartilage tumors, it is unknown how glutamine regulates tumor development in enchondromas, and cancer cell survival in chondrosarcoma. We therefore used a genetically engineered mouse model of enchondromas and primary human chondrosarcoma samples to address these questions. Here we identified that GLS was upregulated in both human patient chondrosarcoma samples with mutations in *IDH1* or *IDH2* and murine chondrocytes with *Idh1* mutation. However, deleting *Gls* in the mouse led to an increased number and size of benign tumor-like lesions and affected hypertrophic chondrocyte differentiation, likely due to a reduction of *α*-KG; whereas inhibiting GLS in *IDH1* or *IDH2* mutant chondrosarcoma led to a smaller tumor size and a reduction in cell viability, associated with the compromised production of non-essential amino acids. Collectively, these data highlight a previously unknown stage-dependent role of glutamine metabolism in cartilage tumors and may provide a therapeutic approach for the malignant cartilage tumor chondrosarcoma

## Results

### *IDH1* or *IDH2* mutant chondrosarcomas cells exhibited increased glutamine contribution to anaplerosis and non-essential amino acids production

From a published dataset showing mRNA profiling (<u>E-MTAB-7264</u>) of chondrosarcoma tumors [40], we observed that expression of *GLS* was upregulated in chondrosarcomas with *IDH1* or *IDH2* mutations (Supplementary Fig 1), indicating glutamine metabolism might be important for cartilage tumor with *IDH1 or IDH2* mutation. To understand how glutamine was utilized in chondrosarcoma cells, we performed carbon tracing experiment with ^13^C5-glutamine in chondrosarcoma cells with wild type *IDH1* and *IDH2*, mutant *IDH1*, and mutant *IDH2*. ^13^C contribution to downstream metabolites was determined by measuring the isotope-labeling pattern. To be utilized by a cell, glutamine is first deaminated to glutamate through GLS. After that, glutamate can be converted to *α*-KG, a key metabolite in the tricarboxylic acid (TCA) cycle (Fig 1A). There was a significant amount of glutamate in chondrosarcomas of different genotypes labeled with ^13^C (Fig 1B). Importantly, the labeling of ^13^C in glutamate was significantly higher in *IDH1* or *IDH2* mutant chondrosarcomas when compared to chondrosarcoma cells with wildtype *IDH1* and *IDH2* (Fig 1B). The ^13^C labeling pattern was examined in the TCA cycle intermediates and non-essential amino acids. Chondrosarcomas with *IDH1* or *IDH2* mutations had significantly higher carbon contribution to all TCA cycle intermediates (Fig 1C-1H) and some non-essential amino acids such as alanine (Fig 1I).

**Fig 1:**
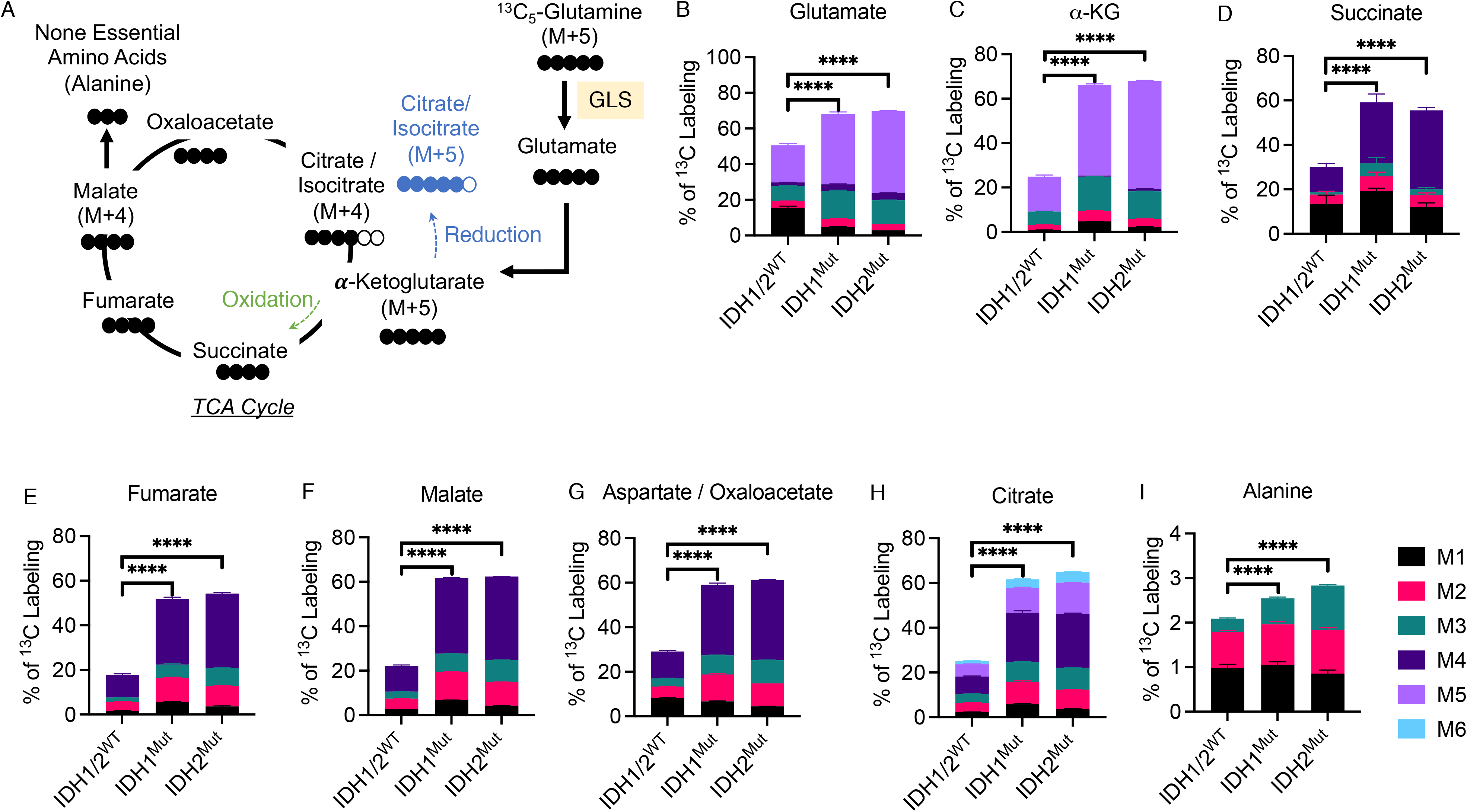
Chondrosarcomas with *IDH1* or *IDH2* mutations had significantly increased contribution from glutamine carbon to downstream metabolites. (A) Graphical depiction of tracing glutamine metabolism using ^5^C5-glutamine. Filled circles indicate ^13^C and open circles indicate ^12^C. Green dashed line with arrowhead indicates the direction of oxidative decarboxylation. Blue dashed line with arrowhead indicates the direction of reductive carboxylation. (B-I) Percentage of ^13^C5-glutamine contribution to glutamate (B), *α*-ketoglutarate (C), succinate (D), fumarate (E), malate (F), oxaloacetate / aspartate (G), citrate (H), and alanine (I). mean±SD are shown. ****p<0.0001 (ANOVA).

Thus, *IDH1* or *IDH2* mutant chondrosarcomas were more efficient than *IDH1* and *IDH2* wildtype chondrosarcomas in converting glutamine to its downstream metabolites.

### GLS was upregulated in IDH mutant chondrocytes

To understand the role of GLS in murine chondrocytes with *Idh1* mutation, we first examined whether expression of mutant *Idh1* gene could lead to altered GLS function. Expression of a mutant IDH1^R132Q^ enzyme in chondrocytes was shown to be sufficient to initiate enchondroma-like lesion formation in mice [41]. Chondrocytes were isolated from mice expressing the conditional IDH1^R132Q^ knock-in allele and transduced with adenovirus GFP or adenovirus Cre [42], GLS activity in these chondrocytes was determined by measuring the conversion from radio-active ^3^H-glutamine to radio-active ^3^H-glutamate. In primary chondrocytes expressing mutant IDH1^R132Q^, GLS activity was significantly upregulated by 2.5-fold (Fig 2A).

**Fig 2:**
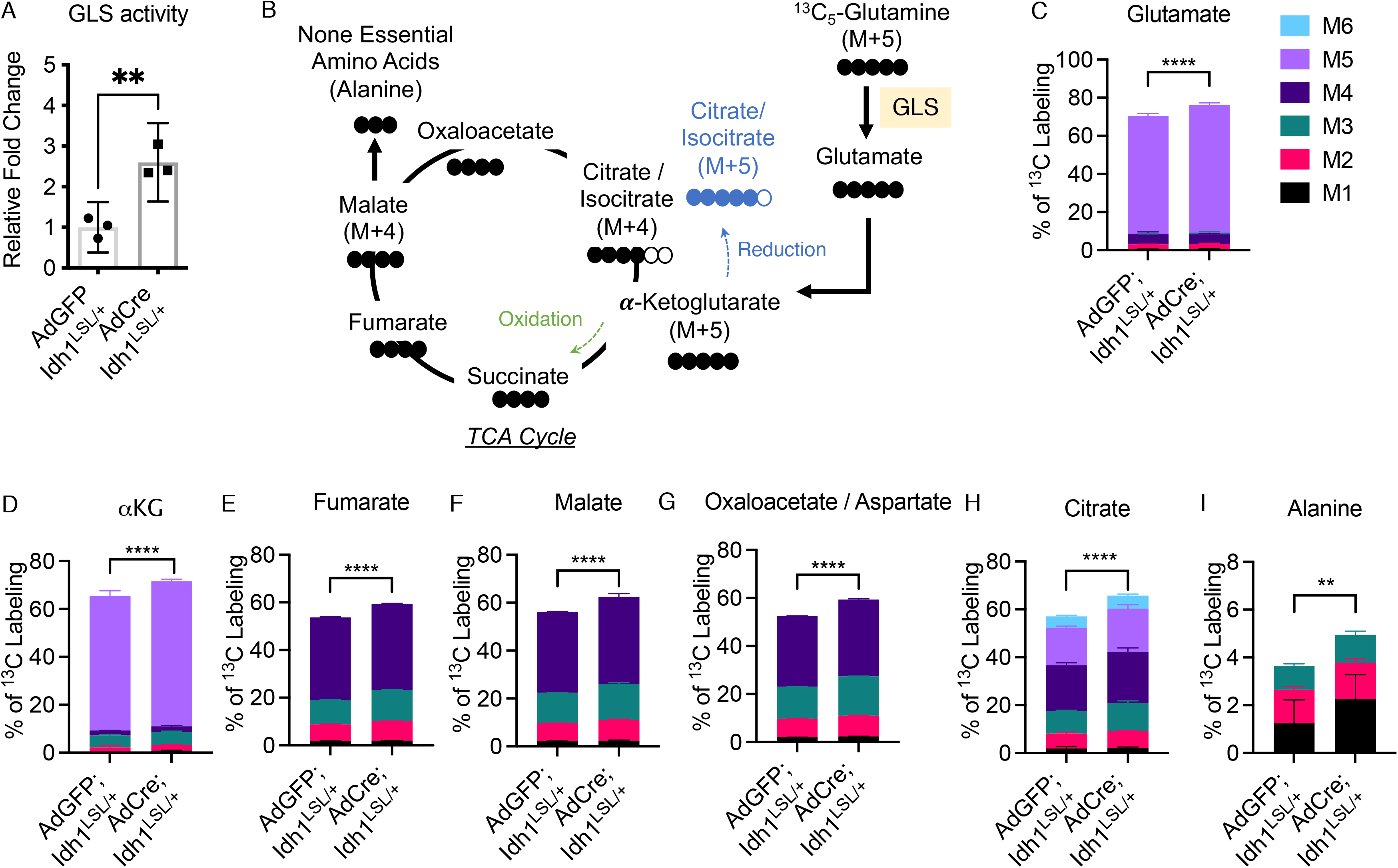
Chondrocytes with *Idh1* mutation had increased GLS activity and glutamine contribution to downstream metabolites. (A) GLS activity of AdGFP;*Idh1^LSL/+^* and AdCre;*Idh1^LSL/+^* chondrocytes. (B) Graphical depiction of tracing glutamine metabolism using ^5^C5-glutamine. Filled circles indicate ^13^C and open circles indicate ^12^C. Green dashed line with arrowhead indicates the direction of oxidative decarboxylation. Blue dashed line with arrowhead indicates the direction of reductive carboxylation. (C-I) Percentage of ^13^C5-glutamine contribution to glutamate (C), *α*-ketoglutarate (D), fumarate (E), malate (F), oxaloacetate / aspartate (G), citrate (H), and alanine (I). M0, M1, …, Mn refers to the isotopologues containing n heavy atoms in a molecule. For (A), mean±95% CI is shown, **p<0.01 (unpaired student t-test). For (C-I), mean±SD are shown. **p<0.01, ****p<0.0001 (ANOVA).

To investigate how chondrocytes with wildtype and mutant IDH1 enzymes utilize glutamine, we conducted ^13^C tracing studies. In chondrocytes expressing the mutant *Idh1*, the percentage of ^13^C labeling in glutamate (Fig 2C), *α*-KG (Fig 2D), other TCA cycle intermediates (Fig 2E-2H), and the non-essential amino acid alanine (Fig 2I) was significantly increased.

Glutamine is a primary source for the production of D-2HG in IDH1 or IDH2 mutant cancers including chondrosarcoma [11, 32, 33]. To examine whether glutamine is also the primary source for D-2HG production in the murine chondrocytes expressing mutant *Idh1*, we cultured these chondrocytes with ^13^C5-glutamine and examined the percentage of ^13^C-labeled D-2HG at different time points. After 10 hours, more than 80% of the D- 2HG was labeled with ^13^C, confirming glutamine is the primary source for D-2HG in IDH1^R132Q^ cells (Supplementary Fig 2). Thus, chondrocytes with an *Idh1* mutation showed increased GLS activity and more efficient conversion of glutamine to downstream metabolites, including D-2HG.

### GLS was important for chondrocyte differentiation and proliferation in chondrocytes with *Idh1* mutation

There are two isoforms of GLS, kidney-type GLS (encoded by *Gls*) and liver-type GLS (encoded by *Gls2*). There was very low level of expression of *Gls2* compared to *Gls* in primary chondrocytes of both *Col2a1*Cre;*Idh1^LSL/+^* and *Col2a1*Cre animals (Fig 3A, GSE123130), consistent with the notion that GLS activity in chondrocytes was mainly catalyzed by *Gls.* We thus focused on *Gls* in the murine chondrocytes. To examine the role of *Gls* in vivo, we conditionally deleted *Gls* in these cells by crossing mice carrying a conditional null allele of *Gls* (*Gls^fl^*) to the *Col2a1*Cre deleter mouse which expresses Cre-recombinase in chondrocytes (Fig 3B, 3C) [43, 44]. *Col2a1*Cre;*Gls^fl/fl^* chondrocytes had reduced glutamine uptake and glutamate production (Fig 3D), confirming the function of GLS was efficiently deleted in these mice.

**Figure 3:**
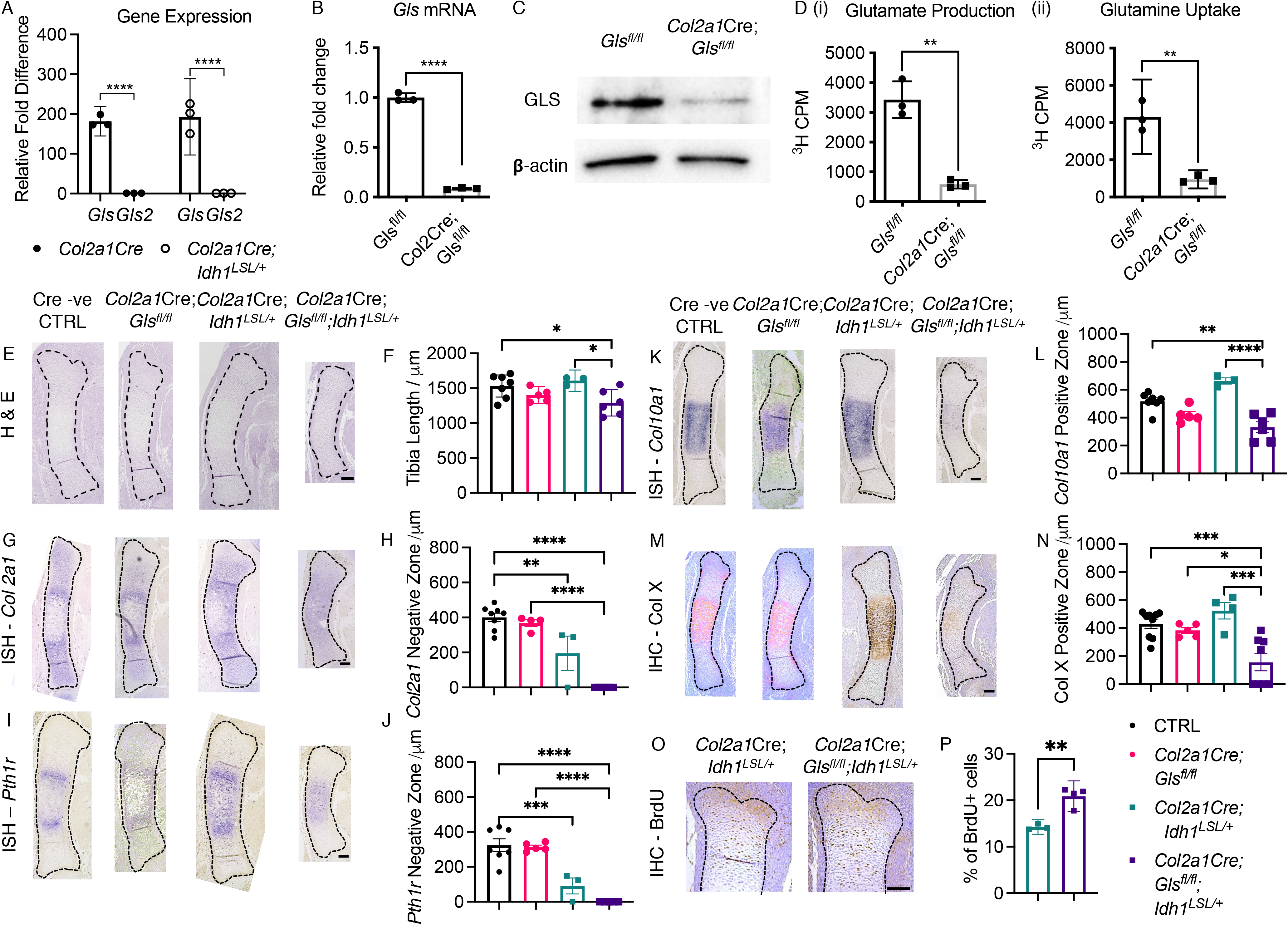
Deleting GLS in *Col2a1*Cre;*Idh1^LSL/+^* chondrocytes affected chondrocyte differentiation. (A) Expression of *Gls* and *Gls2* in chondrocytes of *Col2a1*Cre and *Col2a1*Cre;*Idh1^LSL/+^* animals. (B) Expression of *Gls* in chondrocytes of *Gls^fl/fl^* and *Col2a1*Cre;*Gls^fl/fl^* animals.(C) Western blot of GLS and β;-actin in chondrocytes of *Gls^fl/fl^* and *Col2a1*Cre;*Gls^fl/fl^* animals. (D) ^3^H-glutamate production (i) and ^3^H-glutamine uptake (ii) in chondrocytes of *Gls^fl/fl^* and *Col2a1*Cre;*Gls^fl/fl^* animals. (E) H&E staining. (F) Quantification of the length of tibias. (G) In situ hybridization of *Col2a1*. (H) Quantification of the length of *Col2a1* negative zone. (I) In situ hybridization of *Pth1r*. (J) Quantification of the length of *Pth1r* negative zone. (K) In situ hybridization of *Col10a1.* (L) Quantification of the length of *Col10a1* positive zone. (M) Immunohistochemistry of Col X. (N) Quantification of the length of Col X positive area. (O) Immunohistochemistry of BrdU. (P) Quantification of percentage of BrdU positive cells. For (A), (B), (D), (P), mean±95% CI are shown. **p<0.01, ****p<0.0001 (unpaired student t-test). For (F), (H), (J), (L), (N) mean±SEM are shown. *p<0.05, **p<0.01, ***p<0.001, ****p<0.0001 (ANOVA). Scale bar = 100 μm.

We first examined the phenotype of murine tibias lacking *Gls* in chondrocytes with wildtype or mutant *Idh1* genes using H&E staining (Fig 3E). At embryonic E14.5, deleting *Gls* in *Idh1* wildtype chondrocytes did not cause a significant change in bone length (Fig 3E, 3F). Expression of a mutant *Idh1* gene in chondrocytes at this stage did not cause a reduction in bone length (Fig 3E, 3F). Deleting *Gls* in *Idh1* mutant chondrocytes significantly reduced the bone length (Fig 3E, 3F). At E16.5 and E18.5, we observed reduced bone length and defects in chondrocyte hypertrophic differentiation in *Idh1* mutant mice (Supplementary Fig 3A – 3F), consistent with our previous report [41]. Deleting *Gls* did not cause an obvious skeletal phenotype in *Idh1-* wildtype and *Idh1* mutant mice (Supplementary Fig 3A – 3F).

We then used in situ hybridization to examine markers important for chondrocyte differentiation. Early chondrocyte marker *Col2a1* and hypertrophic chondrocyte marker *Pth1r* were expressed by tibial chondrocytes at E14.5 (FIg 3G – 3J). However, the region between *Col2a1* expressing cells and *Pth1r* expressing cells were reduced in *Idh1* mutant animals. In *Gls;Idh1* double mutant mice, *Col2a1* expression cells and *Pth1r* expressing cells became single regions in the middle of the bone with no separation (Fig 3G-3J). *Col10a1* is expressed by hypertrophic chondrocytes. At E14.5, expression pattern of *Col10a1* was comparable among control, *Gls* mutant, and *Idh* mutant mice, but in *Gls;Idh1* double mutant mice there was reduced zone of staining (Fig 3K – 3N). We also examined proliferation and apoptosis in these animals. No difference in proliferation was determined between control and *Gls* mutant animals in *Idh1* wildtype background, but there was an increase in proliferation when *Gls* was deleted in *Idh1* mutant mice (Fig 3O, 3P). Apoptosis was not detected in all animals at this stage based on immunohistochemistry of cleaved caspase 3 (Supplementary Fig 3G).

### Deleting GLS in chondrocytes increased the number and size of enchondroma- like lesions

We next examined how deleting *Gls* in chondrocyte postnatally affected chondrocyte homeostasis in growth plates and enchondroma-like lesion formation in adult animals. We induced deletion of *Gls* at four weeks of age by tamoxifen and examined the phenotype at 6 months of age, a time point when growth plates were completely remodeled in control animals and enchondroma-like lesions were stable in *Idh1* mutant mice. Enchondroma-like lesions were identified in *Idh1* mutant animals as we previously reported and these animals showed less trabecular bone volume (Fig 4A - 4D). No obvious growth plate phenotype was observed in *Gls* mutant mice and histologic examination showed less trabecular bone (Fig 4A, 4B). In contrast, the number and size of enchondroma-like lesions were significantly increased when *Gls* was deleted in *Idh1* mutant animals, and the trabecular bone volume was further reduced (Fig 4A – 4D).

**Fig 4:**
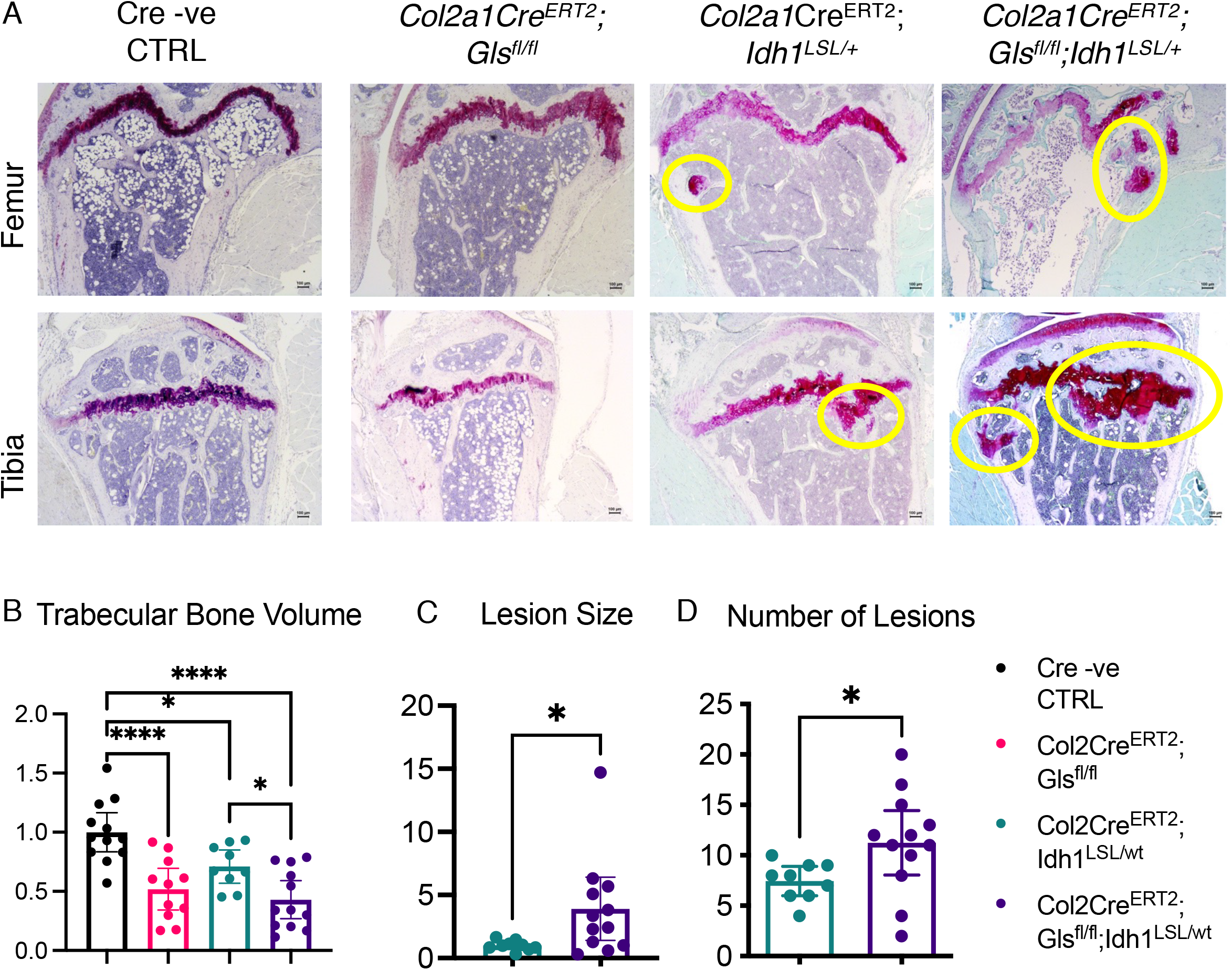
Deleting *Gls* in *Col2a1*Cre^ERT2^;*Idh1^LSL/+^* chondrocytes increased the number and size of enchondroma-like lesions. (A) Representative Safranin O staining of mice of specified genotypes. Enchondroma-like lesions are highlighted in yellow circles. (B) Quantification of the volume of trabecular bone based on Safranin O staining. (C-D) Quantification of the number (B) and size (C) of enchondroma-like lesions in animals of specified genotypes. mean±95% CI are shown, For (B), mean±SEM are shown. *p<0.05, ****p<0.0001 (ANOVA). For (C) and (D), *p<0.05 (unpaired student t-test). Scale bar = 100 μm.

### GLS regulated chondrocyte differentiation through *α*-ketoglutarate

Glutamine metabolism regulates chondrocyte differentiation through its downstream metabolites *α*-KG in *Idh1* wildtype background [28]. We examined whether GLS could regulate the differentiation of chondrocytes containing a *Idh1* mutation through *α*-KG. Our previous ^13^C tracing experiments showed *α*-KG was mainly derived from glutamine and *Idh1* mutant chondrocytes were more efficient in converting glutamine to *α*-KG (Fig 2D). We found the level of *α*-KG was reduced by more than 70% in *Idh1* mutant chondrocytes (Fig 5A). Because chondrocytes with *Idh1* mutation were more efficient in converting glutamine to *α*-KG, the lower intracellular concentration was likely because *α*-KG was converted to D-2HG. When GLS was inhibited in *Idh1* mutant chondrocytes by Bis-2-(5-phenylacetamido-1,3,4-thiadiazol-2-yl)ethyl sulfide (BPTES), a small molecular inhibitor for GLS [45], *α*-KG level was further reduced by about 60% (Fig 5B).

**Fig 5:**
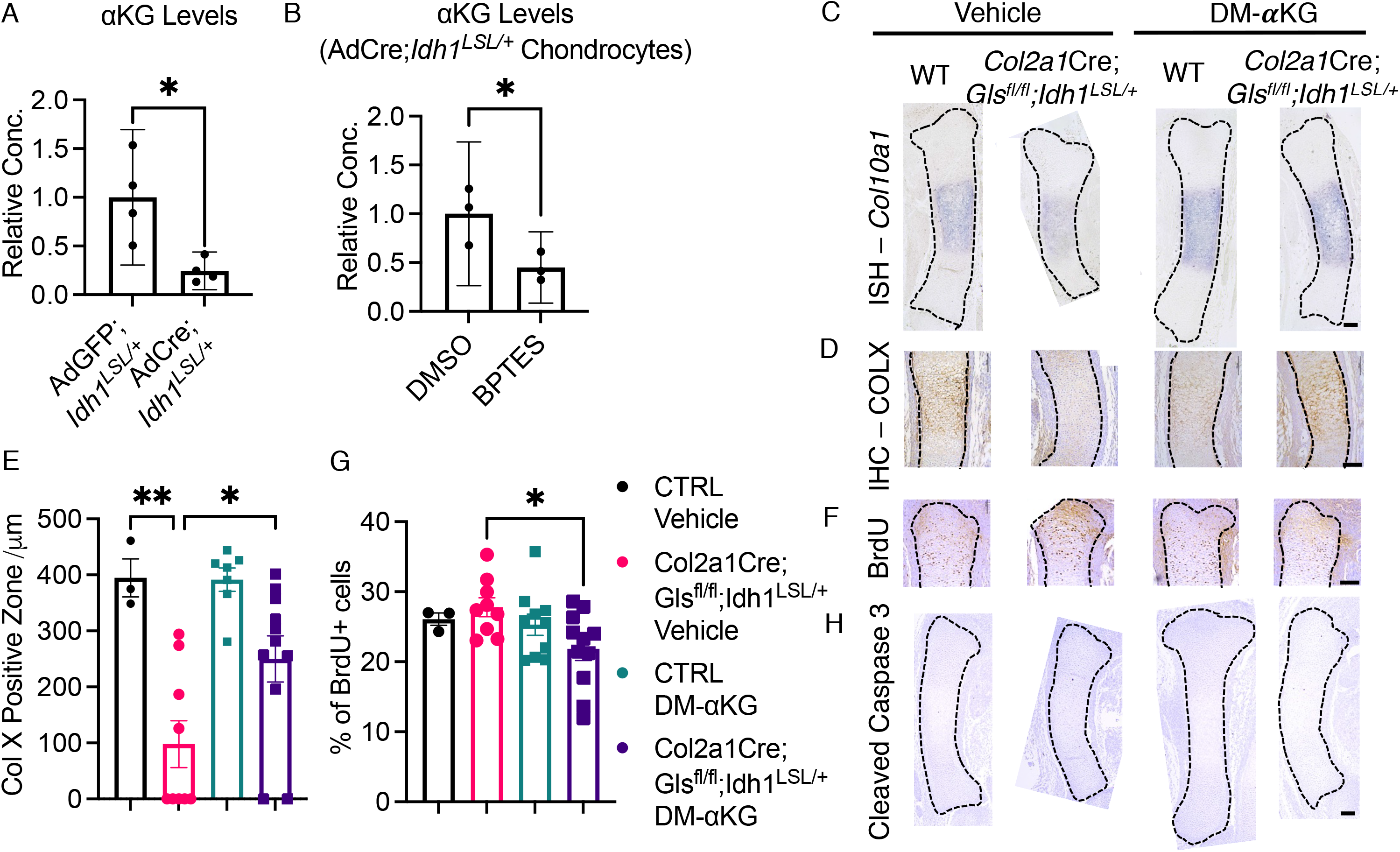
GLS regulated *Col2a1*Cre;*Idh1^LSL/+^* chondrocyte differentiation through downstream metabolite *α*-KG. (A) Relative intracellular *α*-KG concentration in AdGFP;*Idh1^LSL/+^* and AdCre;*Idh1^LSL/+^* chondrocytes. (B) Relative intracellular *α*-KG concentration in AdCre;*Idh1^LSL/+^* chondrocytes treated with vehicle control or 10 μM BPTES. (B) Immunohistochemistry of Col X of metatarsal bones in organ culture. (C) In situ hybridization of *Col10a1.* (D) Immunohistochemistry of Col X. (E) Quantification of the length of Col X positive zone. (F) Immunohistochemistry of BrdU. (G) Quantification of the percentage of BrdU positive cells. (H) Immunohistochemistry of cleaved caspase 3. For (A), (B), mean±95% CI are shown. *p<0.05 (unpaired student t-test). For (E), (G), mean±SEM are shown. *p<0.05, **p<0.01(ANOVA). Scale bar = 100 μm.

Next, we investigated whether exogenous *α*-KG could rescue the defects of chondrocyte differentiation in *Col2a1*Cre;*Gls^fl/fl^*;*Idh1^LSL/+^* animals. We injected cell permeable dimethyl-*α*-KG (DM-*α*-KG) to pregnant dams every day from E11.5 to E13.5 and harvested the embryos at E14.5. Treatment with DM-*α*-KG rescued chondrocyte hypertrophy as *Col10a1* mRNA and COLX protein expression were both observed in *Gls;Idh1*- double mutant mice (Fig 5C – 5E). Finally, DM-*α*-KG treatment led to a modest decrease in proliferation (Fig 5F, 5G) without inducing apoptosis (Fig 5H).

### Inhibiting GLS in *IDH1* or *IDH2* mutant chondrosarcoma xenografts reduced tumor weight

Previously it was reported that inhibiting GLS reduced viability in chondrosarcoma cell lines [36]. However, our murine data in enchondroma-like lesions suggests that GLS inhibition would have an opposite effect. We thus established patient-derived xenograft models as previously described to examine the effects of GLS inhibition in chondrosarcoma [46]. Chondrosarcoma tumors were divided into pieces at 5 mm x 5 mm x 5 mm and implanted to immune-deficient Nod-scid-gamma mice subcutaneously. We treated these animals with BPTES or vehicle control for 14 days and measured the tumor weight at the time of sacrifice. For each patient sample, the tumor weight of each mouse was normalized to the average tumor weight in the vehicle control group of the same patient sample. BPTES treatment significantly reduced tumor weight of chondrosarcoma tumors with *IDH1* mutations (Fig 6B, 6C). We examined proliferation and apoptosis in these tumors using immunohistochemistry of Ki67 and cleaved caspase 3 respectively. BPTES treatment significantly increased apoptosis in chondrosarcoma xenografts but we did not observe a substantial difference in proliferation (Fig 6D-6G). We then examined chondrosarcoma cells treated with BEPTS in vitro. Consistent with our findings in vivo, BPTES treatment reduced cell viability, increased cell apoptosis, and did not alter proliferation in *IDH1* or *IDH2* mutant chondrosarcomas (Fig 6H-6J). Interestingly, BPTES treatment did not alter tumor burden of *IDH1* and *IDH2* wildtype chondrosarcoma (Supplementary Fig 4). These data support previously published studies and show a different role of GLS in human chondrosarcoma than murine enchondroma-like lesions.

**Fig 6:**
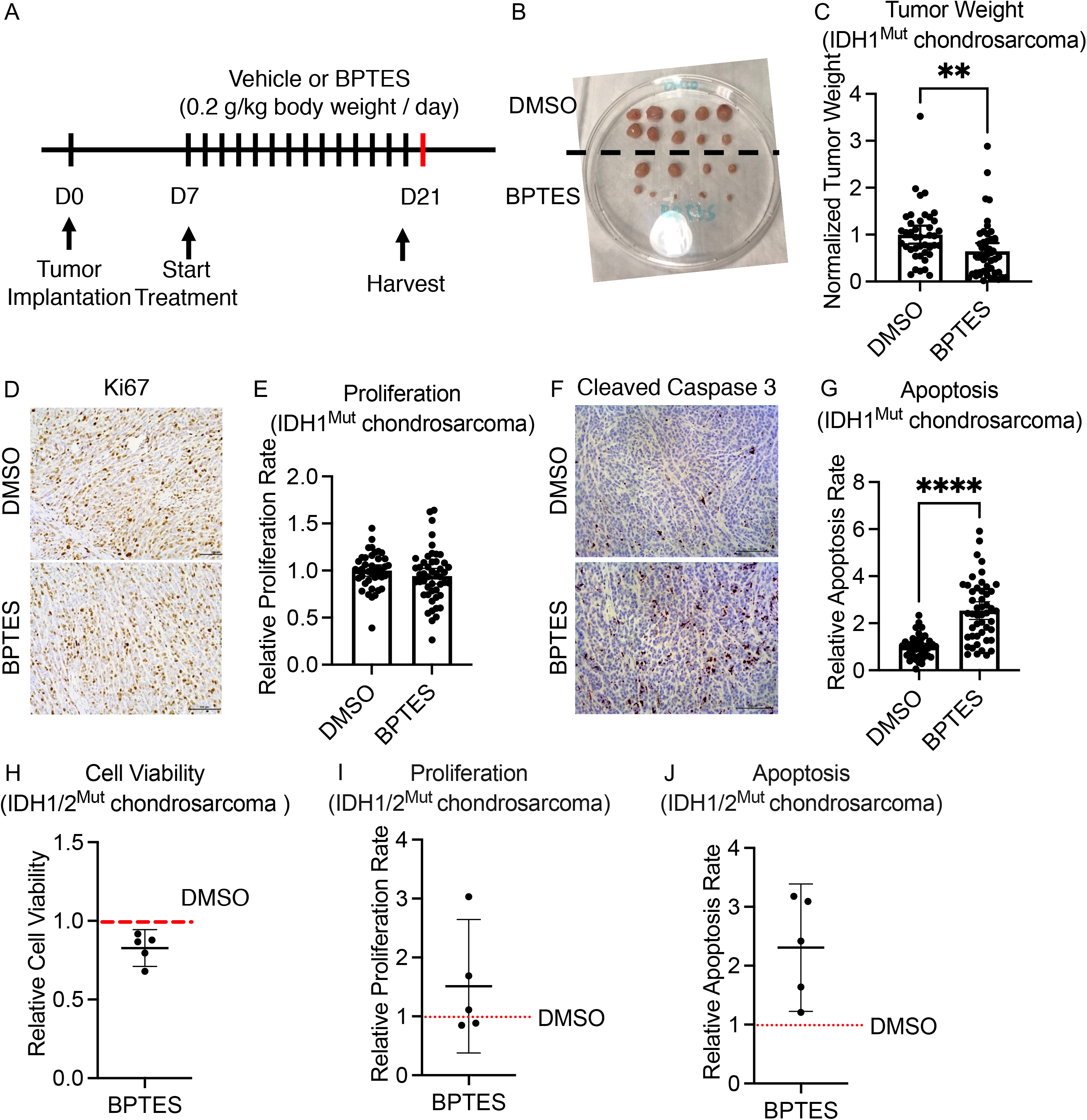
Inhibiting GLS reduced tumor weight and induced apoptosis in chondrosarcomas with *IDH* mutations. (A) Experimental design for the xenograft experiment. (B) Representative pictures of xenografted tumor at the time of harvest. (C) Relative tumor weight of xenografted chondrosarcoma tumors at the time of harvest. (D) Representative picture of immunohistochemistry of Ki67 of xenografted tumors. (E) Quantification of relative proliferation rate of each tumor determined by percentage of Ki67 positive cells. (F) Representative picture of immunohistochemistry of cleaved caspase 3 of xenografted tumors. (G) Relative apoptotic rate of each tumor determined by percentage of cleaved caspase 3 positive cells. (H) Relative cell viability of IDH1/2^Mut^ chondrosarcoma cells treated with 10μM BPTES determined by CellTiter Glo cell viability assay. (I) Relative proliferation rate of *IDH1* or *IDH2* mutant chondrosarcoma cells treated with 10μM BPTES in vitro determined by EdU staining. (J) Relative apoptosis of *IDH1* or *IDH2* mutant chondrosarcoma cells treated with 10μM BPTES in vitro determined by TUNEL staining. For 6H-6J, Each dot represents one patient sample. mean±95% CI are shown. **p<0.01. ****p<0.0001 (unpaired student t-test).

### GLS inhibited the production of non-essential amino acids in chondrosarcoma

We next sought to understand how glutamine metabolism regulated cell survival of *IDH1* or *IDH2* mutant chondrosarcomas. Similar to in the mouse, *α*-KG levels were lower with BPTES treatment (Fig 7A), However, unlike our murine findings, adding DM-*α*-KG did not rescue the apoptosis changes seen with BEPTS treatment (Fig 7B).

**Fig 7:**
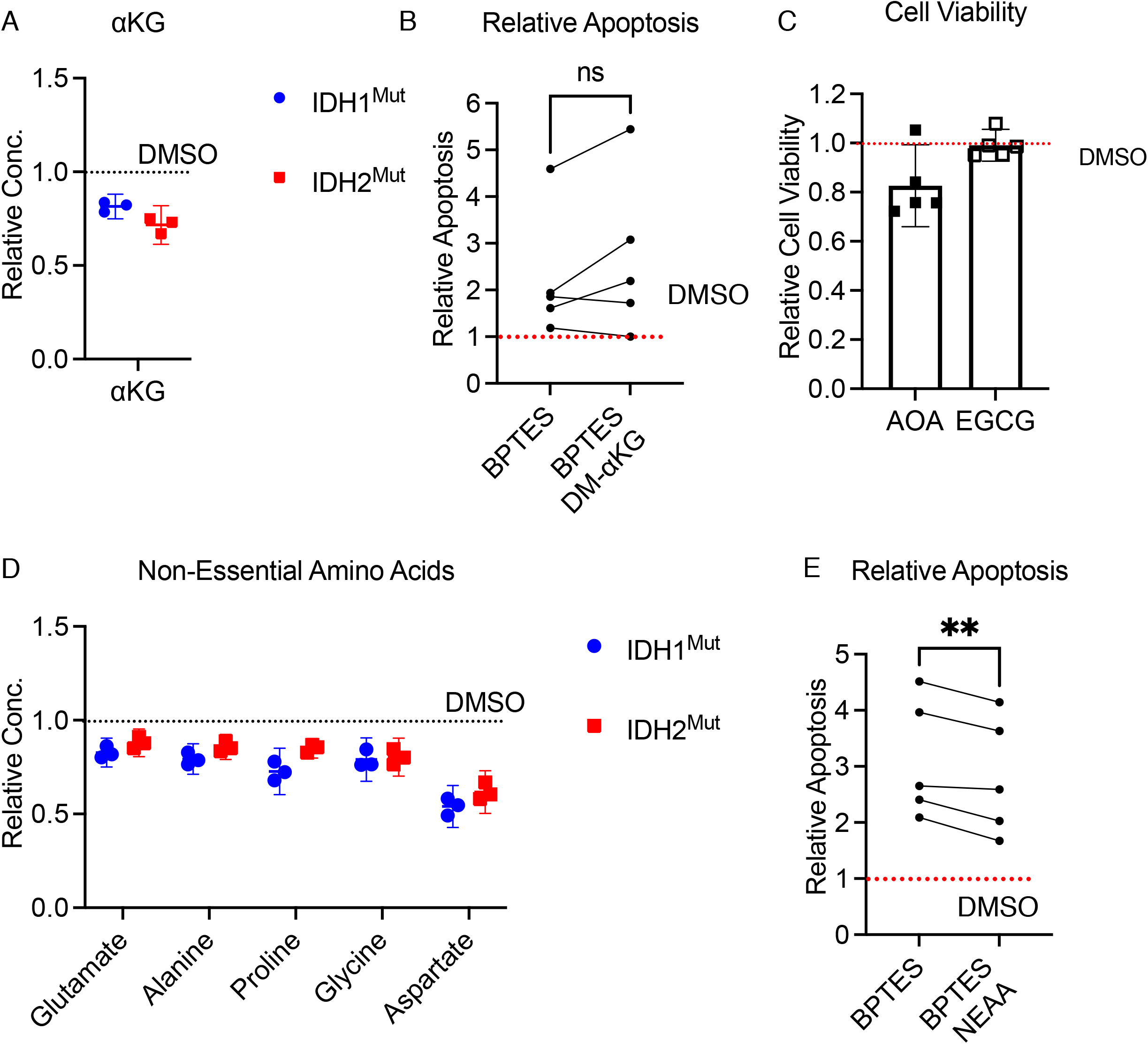
GLS regulated cell apoptosis of chondrosarcomas with *IDH1* or *IDH2* mutations through production of non-essential amino acids. (A) Relative concentration of *α*-KG in *IDH1* or *IDH2* mutant chondrosarcomas treated with 10μM BPTES. (B) Relative apoptosis of *IDH1* or *IDH2* mutant chondrosarcoma cells treated 10μM BPTES, or 10μM BPTES+1mM DM-*α*-ketoglutarate determined by Annexin V staining. (C) Relative cell viability of IDH1/2^Mut^ chondrosarcoma cells treated with 100μM AOA and and 500μM EGCG. Each dot represents one patient sample. (D) Relative concentration of different amino acids in *IDH1* or *IDH2* mutant chondrosarcomas treated with 10μM BPTES. (E) Relative apoptosis of *IDH1* or *IDH2* mutant chondrosarcoma cells treated 10μM BPTES, or 10μM BPTES+2X NEAA determined by Annexin V staining. For 7(B) and 7(E), each dot represents one patient sample. Relative metabolite concentration (7A, 7D), apoptosis (7B, 7E), and cell viability (7C) were determined by normalizing the value of each treatment group to the value of the vehicle control in each experiment. For (A), (C), and (D), mean±95% CI are shown. For (B) and (E), **p<0.01 (paired student t- test).

Downstream of GLS, glutamate could be further metabolized to *α*-KG through glutamate dehydrogenase and transaminases. Inhibiting transaminases by aminooxyacetate (AOA) caused a reduction of cell viability in *IDH1* or *IDH2* mutant chondrosarcomas similar to the effects of BPTES, but inhibiting glutamate dehydrogenase by epigallocatechin gallate (EGCG) did not affect cell survival (Fig 7C). These data suggested transaminases might be more critical for chondrosarcoma cell survival downstream of GLS.

Non-essential amino acids can be produced from glutamine through GLS and then transaminases. They are known to play role in regulating cancer apoptosis, raising the possibility that they are playing this role in chondrosarcoma. We found lower levels of multiple non-essential amino acids when GLS was inhibited by BPTES (Fig 7D). We treated these chondrosarcoma cells with non-essential amino acids, and in contrast to our data from DM-*α*-KG, we found a modest rescue of apoptosis changes (Fig 7E). However, supplementing non-essential amino acids to *Col2a1*Cre;*Gls^fl/fl^*;*Idh1^LSL/+^* metatarsal organ culture did not lead to rescue of chondrocyte differentiation defects (Supplementary Fig 5), suggesting they might not be the key mediator of the murine enchondroma-like lesions.

## Discussion

In this study, we found that glutamine contribution to downstream metabolites was upregulated in chondrosarcomas and chondrocytes with mutations in IDH1 or IDH2 enzymes. Glutamine metabolism played different roles in the benign tumor enchondroma and the malignant cancer chondrosarcoma. Deleting GLS in murine chondrocytes with *Idh1* mutation interrupted hypertrophic differentiation during embryonic development and led to increased number and size of enchondroma-like lesions in adult animals. In malignant *IDH1* or *IDH2* mutant chondrosarcomas isolated from human patients, pharmacological inhibition of GLS led to reduced tumor weight.

Enchondroma arises from dysregulated chondrocyte differentiation. In our mouse data, we observed that chondrocytes with an *Idh1* mutation had significantly lower *α*-KG levels and blocking GLS further decreased *α*-KG concentration. It is possible that glutamine metabolism regulates chondrocyte differentiation and the enchondroma-like phenotype of *Idh1* mutant animals through its downstream metabolites *α*-KG. Glutamine metabolism is known to regulate cell proliferation and differentiation in the skeletal system. Stegen et al. reported *Gls* regulated chondrocyte differentiation through *α*-KG and its downstream metabolite acetyl-CoA postnatally via epigenetic regulation [28]. Yu et al. showed deleting *Gls* in skeletal stem cells led to increased adipogenic differentiation and compromised osteogenic differentiation as well as reduced cell proliferation [27]. Our finding is consistent with these studies and further support the notion that *Gls* could regulate cell differentiation and proliferation through *α*-KG in the skeletal system. The mechanism through which *α*-KG regulate chondrocyte differentiation remains to be determined. It possible that *α*-KG could affect expression of chondrogenic genes through altering histone modifications and DNA methylation.

Different from our data in mouse enchondroma-like lesions, inhibiting GLS in human *IDH1* or *IDH2* mutant chondrosarcomas led to reduced tumor weight. Supplementing non-essential amino acids to BPTES treated *IDH1* or *IDH2* mutant chondrosarcomas partially rescued apoptosis, suggesting these cancer cells relied on glutamine metabolism for the production of non-essential amino acids to prevent cell death. In multiple different cancer cell lines, glutamine supported cancer cell proliferation and suppressed apoptosis and autophagy through producing other non-essential amino acids [47–51]. Non-essential amino acids mixture provides seven non-essential amino acids alanine, asparagine, aspartate, glycine, serine, proline, and glutamate. Many of them have been shown to be critical for cancer cell viability, especially under metabolic stress. In glioblastoma and neuroblastoma, glutamine deprivation induced apoptosis, which could be restored by exogenous asparagine [51]. In pancreatic cancer, aspartate becomes “essential” when glutamine availability or metabolism is limited [52]. Glutamine could also regulate cancer cell viability through sustaining cellular redox homeostasis [26]. For the next step, it will be important to understand which and how non-essential amino acid(s) prevent apoptosis in *IDH1* or *IDH2* mutant chondrosarcomas.

The distinct roles of glutamine metabolism in enchondroma and chondrosarcoma were possibly due to different metabolic needs at different stages of tumor development. As enchondroma rises from dysregulated chondrocyte differentiation, *α*-KG – the regulator for cell differentiation plays an important role in the development of the benign tumor. For malignant chondrosarcomas, changes in differentiation might not be critical for tumor growth. However, these *IDH1* or *IDH2* mutant cancer cells might require some amino acids to support anaplerosis or synthesize glutathione in order to cope with oxidative stress.

One limitation of this study is that we studied the role of glutamine metabolism in enchondromas and chondrosarcomas using mouse models and human patient samples respectively. Thus, it is possible the distinct roles of glutamine metabolism we observed was caused by different metabolic needs of murine and human cells. To address this concern, it will be important to establish mouse models to study chondrosarcoma and to determine the role of glutamine metabolism in human enchondroma samples.

In summary, our study showed that GLS-mediated glutamine metabolism played distinct roles in *IDH1* or *IDH2* mutant enchondromas and chondrosarcomas although it was upregulated in both conditions. In the context of enchondroma, deleting GLS in chondrocytes with *Idh1* mutation increased the number and size of enchondroma-like lesions. In chondrosarcoma, inhibiting GLS led to decreased tumor weight. Glutamine regulated the proliferation and apoptosis in these tumors differently through different downstream metabolites. Glutamine-derived *α*-ketoglutarate is a key regulator of chondrocyte differentiation. Deleting GLS in chondrocytes with *Idh1* mutation impaired hypertrophic differentiation and increased cell proliferation, which may lead to increased number and size of enchondroma-like lesions. In chondrosarcoma, GLS regulated cell apoptosis partially through producing non-essential amino acids. Inhibiting GLS reduced cell viability and increased cell apoptosis. Supplementation of non-essential amino acids partially rescued such effects.

## Methods

### Animals

*Idh1^LSL/+^*, *Gls^fl/fl^* (JAX: #017894), *Col2a1*Cre , *Col2a1*Cre^ERT2^, and NOD *scid* gamma (NSG) mice are as previously described [27, 41, 43, 44, 53, 54]. Congenic animals were used in every experiment. *Idh1^LSL/+^* mice bear a conditional knock-in of the point mutation IDH1-R132Q as previous clarified [41].

### Isolation of primary chondrocytes

Primary sternal chondrocytes isolation was as described before [55]. In brief, mouse sterna and ribs from P3 neonates were cleaned and digested by pronase (Roche, 11459643001) at 2 mg/ml PBS at 37°C with constant agitation for 1 hour, Collagenase IV (Worthington, LS004189) at 3 mg/ml DMEM at 37°C for 1 hour, Collagenase IV at 0.5 mg/ml DMEM at 37°C for 3 hours, and filtered using 45 μm cell strainer.

### Cell Culture

Chondrocytes were cultured in high-glucose DMEM with 10% FBS and 1% penicillin / streptomycin. Primary chondrosarcoma cells were cultured in *α*-MEM with 10% FBS and 1% penicillin / streptomycin. In some experiments, chondrocytes and chondrosarcoma cells were treated with 10μM Bis-2-(5-phenylacetamido-1,2,4-thiadiazol-2-yl)ethyl sulfide (BPTES), 100μM epigallocatechin gallate (EGCG), 500μM aminooxyacetate (AOA), 2X non-essential amino acids, or DMSO at indicated for indicated 48 hours. Cells were cultured at 37°C humidified 5% CO2 incubator.

### Transfection

In indicated experiment, chondrocytes were transfected with adenovirus-GFP and adenovirus-Cre at 400 MOI.

### Metatarsal Organ Culture

The 2^nd^, 3^rd^, and 4^th^ metatarsal bones were dissected from the hindlimbs of embryos at E16.5. They were transferred to 24-well non-adherent plate with 1mL α-MEM supplemented with 50μg/ml ascorbic acid, 1mM β-glycerophosphate, and 0.2% bovine serum albumin. The media was changed every other day. Explants were cultured at 37°C humidified 5% CO2 incubator for five days and then fixed with 10% neutral buffered formalin (NBF).

### EdU assay

EdU assay (ThermoFisher C10337) was performed according to manufacturer’s instructions. In brief, cells were cultured with 10 mM EdU for 12 hours prior to fixation with 4% PFA / PBS for 15 min and permeabilization with 0.5% Triton*™* X-100 for 20 min at room temperature. Cells were then incubated with Click-iT® reaction cocktail for 30 min in dark at room temperature and stained with DAPI.

### TUNEL assay

TUNEL assay was performed according to manufacturer’s instructions (Roche, 11684795910). In brief, cells were fixed with 4% PFA / PBS at room temperature for 1 hr and permeabilized with 0.1% Triton*™* X-100 in 0.1% sodium citrate for 2 min on ice. After rinse with PBS, cells were incubated with TUNEL reaction mixture at 37°C for 1 hr and stained with DAPI.

### Cell Viability Assay

Cell viability was determined by CellTiter-Glo® Assay according to manufacturer’s instructions (Promega G7570). In brief, CellTiter-Glo® Buffer was thawed at room temperature and transferred to CellTiter-Glo® Substrate. Cell culture plate was equilibrated to room temperature for 30min. 100 μL of CellTiter-Glo® Reagent was added to the cell culture media in each well. The plate was mixed for 2 min on an orbital shaker and incubated for 10 minutes at room temperature. Luminescence was then recorded. Cell viability of each primary chondrosarcoma patient sample was normalized to the average cell viability in the vehicle group of the same sample.

### Annexin V / PI staining

Annexin V and Propidium Iodide staining was performed according to manufacturer’s instructions (Invitrogen™ R37176, Invitrogen™P1304MP). In brief, 1 drop of Annexin V APC Ready Flow Conjugate was added to 0.5 mL of annexin-binding buffer with 2.5 mM calcium. 1mg/mL PI was added to the APC binding buffer with 1/1000 dilution. The cells were incubated for 15 minutes at room temperature. Fluorescence was detected by flow cytometry.

### GLS activity assay

Primary chondrocytes were cultured to prior to experiment. After washing cells with Hanks Buffered Saline Solution (HBSS) for three times, cells were cultured in *α*-MEM containing media containing 2μM Glutamine and 4 mCi/mL L-[3,4-^3^H(N)]-Glutamine (PerkinElmer, NET551250UC). GLS activity was terminated by washing cells with ice cold HBSS for three times followed by scaping cells with 1 mL ice cold milliQ water. Cells were lysed by sonication for 1 min with 1 sec pulse at 20% amplitude. Cell lysates were bound onto AG 1-X8 polyprep anion exchange column. Uncharged glutamine was eluted with 2 mL of milliQ water for three times. Negatively charged glutamate and downstream metabolites were eluted with 2 mL of 0.1M HCl for three times. After adding 4 mL of scintillation cocktail to the eluent, DPM of the solution was measured by a Beckman LS6500 Scintillation counter.

### Xenograft

Prior to the experiment, chondrosarcoma cells from each patient’s tumor were maintained subcutaneously in vivo in NSG mice. For the xenograft experiment, tumors were surgically removed from each mouse and divided into explants of 5 x 5 x 5 mm each, and implanted into the subcutaneous tissue on the pack of NSG mice. BPTES and vehicle control (10% DMSO / PBS) treatment started 10 days after implantation. Mice were treated with BPTES at 0.2 g / kg or vehicle via intraperitoneal injection daily for 14 days. Tumor weights were recorded upon harvest. For each patient-derived xenograft experiment, relative tumor weight was determined by normalizing the tumor weight of each tumor to the average tumor weight of the vehicle control group.

### Histological analysis

Bone histomorphometry for adult mice was performed on hindlimbs fixed in 10% neutral buffered formalin for 3 days followed by decalcification with 14% EDTA for 2 weeks at room temperature. Histomorphometry for embryonic skeletons was performed on tibias fixed in 10% neutral buffered formalin overnight followed by decalcification with 14% EDTA overnight at room temperature. Following decalcification, skeletons were embedded in paraffin and sectioned at 5 μm thickness. Safranin O staining was performed following standard protocol.

### Immunohistochemistry

Immunohistochemistry was performed on 5 μm paraffin-sectioned limbs. For type X collagen, antigen retrieval was performed by citrate buffer incubation at 85°C for 15min and hyaluronidase digestion at 10 mg/ml at 37°C for 30 min. For BrdU staining, BrdU labeling reagent (Invitrogen, 000103) was injected to pregnant female mice at 1 mL / 100 g body weight 2 hours prior to euthanasia. Antigen retrieval was performed by proteinase K digestion at 10 μg/ml at room temperature for 10 min. For MMP13, antigen retrieval was performed by hyaluronidase digestion at 5 mg/ml at 37°C for 30 min. For all immunohistochemistry, endogenous peroxidase activity was blocked by incubation with 3% H2O2 / Methanol for 10 minutes followed by incubation with Dako Dual Endogenous Enzyme Block reagent (Agilent Dako, S2003) for 30 min at room temperature. The specimen was blocked with 2% horse serum at room temperature for 30 min followed by incubation with antibodies for Col X (1:500, ThermoFisher, 14-9771- 82), BrdU (1:1000, ThermoFisher, MA3-071), MMP13 (1:100, MilliporeSigma MAB13424) overnight at 4°C. TUNEL assay was performed according to manufacturer’s instructions (Roche, 11684795910). In brief, the tissue was incubated with proteinase K at 10 μg/ml at room temperature for 20 min, and then in TUNEL labeling reagent 37°C for 1 hour. Quantification of the length of hypertrophic zone and bone elements was done using the image processing software Fiji Image J.

### In situ hybridization

In situ hybridization was performed on 5 μm paraffin-sectioned limbs. Paraffin sections were deparaffinized and rehydrated, followed by fixation with 4% PFA at room temperature for 15 minutes. Sections were then treated with 20 μg/ml proteinase K for 15 minutes at room temperature, fixed with 4% PFA at room temperature for 10 minutes, and acetylated with for 10 minutes. Sections were then incubated with hybridization buffer at 58°C for 3 hours, and then incubated with Digoxigenin-labeled RNA probe (*Col2a1, Pth1r, Co10a1*) at 58°C overnight. Sections were then washed with 5x SSC for one time at 65°C, followed by RNAse A treatment at 37°C for 30 minutes. Sections were then washed 2x SSC for one times, 0.2x SSC for two times at 65°C, and then blocked with 2% Boehringer Blocking Reagent / 20% Heat Inactivated Sheep Serum for one hour. Sections were then blocked with Anti-Digoxigenin antibody (1:4000 in blocking solution) at 4°C overnight. Sections were developed with BM Purple at room temperature until color developed.

### Analysis of enchondroma-like lesions and trabecular bone volume

Tamoxifen was administered daily for 10 days at 100 mg / kg body weight / day via intraperitoneal injection starting at 4 weeks of age. Mice were euthanized at 6-month of age. Hindlimbs were harvested for analyzing growth-plate and enchondroma-like phenotype. Hip-joint cartilage was used for confirming DNA recombination.

Quantification of enchondroma-like lesions was performed as previously described [55]. In brief, enchondroma-like lesions were first identified by Safranin-O staining, which was performed on one slide (2 sections, 5 μm / section) in every ten consecutive slides (10 μm). We then examined every slide consecutively under the light microscope to identify lesions that did not span to the Safranin O stained slide. The number of lesions were then recorded. For every Safranin O stained slide, we manually outlined each lesion and measured the area of each lesion using the image processing software Fiji Image J. We estimated the lesion size of each animal by adding up the areas of all the lesions in that animal. The lesion size for each animal was normalized to the average lesion size of *Col2*Cre^ERT2^;*Idh1^LSL/+^* animals. Quantification of trabecular bone volume was performed by adding up the trabecular bone surface of each Safranin O-stained slide. For every Safranin O stained slide, we manually outlined the trabecular bone 1200 μm below the growth plates of each tibia and femur and measured the area of trabecular bones using the image processing software Fiji Image J. We estimated the lesion size of each animal by adding up the areas of all the trabecular bones in our region of interest in that animal.

### Xenograft

Prior to the experiment, chondrosarcoma cells from each patient’s tumor were maintained subcutaneously in vivo in NSG mice. For the xenograft experiment, tumors were surgically removed from each mouse and divided into explants of 5 x 5 x 5 mm each, and implanted into the subcutaneous tissue on the pack of NSG mice. BPTES and vehicle control treatment started 10 days after implantation. Mice were treated with BPTES at 0.2 g / kg or vehicle control via intraperitoneal injection daily for 14 days.

Tumor weights were recorded upon harvest. For each patient-derived xenograft experiment, relative tumor weight was determined by normalizing the tumor weight of each tumor to the average tumor weight of the vehicle control group.

### D-2HG measurement

D-2HG is analyzed by liquid chromatography – tandem mass spectrometry (LC/MS/MS). Cell culture media was collected from each sample. After adding 2-HG-^2^H4 (internal standard), the sample was dried under nitrogen and derivatized by (+)-O,O’- diacetyl-L-tartaric anhydride (DATAN) for measurement.

### Carbon Isotope Labeling

Chondrocytes were cultured in DMEM with 4500mg/L Glucose and 4mM ^15^C5- Glutamine in 6cm cell culture plates for specified time. 500μL methanol was used to extract metabolites from each plate. After centrifuge at 12000rpm for 15min, supernatant was dried at 37℃. The dried residues were resuspended in 25 μL of methoxylamine hydrochloride (2% (w/v) in pyridine) and incubated at 40°C for 1.5 hours in a heating block. After brief centrifugation, 35 μL of MTBSTFA + 1% TBDMS was added, and the samples were incubated at 60°C for 30 minutes. The derivatized samples were centrifuged for 5 minutes at 20,000 x g, and the supernatants were transferred to GC vials for GC-MS analysis. A modified GC-MS method was employed ^23^. The injection volume was 1 μL, and samples were injected in splitless mode. GC oven temperature was held at 80°C for two minutes, increased to 280°C at 7°C/min, and held at 280°C for a total run time of forty minutes. GC-MS analysis was performed on an Agilent 7890B GC system equipped with a HP-5MS capillary column (30 m, 0.25 mm i.d., 0.25 µm-phase thickness; Agilent J&W Scientific, Santa Clara, CA), connected to an Agilent 5977A Mass Spectrometer operating under ionization by electron impact (EI) at 70 eV. Helium flow was maintained at 1 mL/min. The source temperature was maintained at 230°C, the MS quad temperature at 150°C, the interface temperature at 280°C, and the inlet temperature at 250°C. Mass spectra were recorded in mass scan mode with m/z from 50 to 700.

### 13C-based Stable Isotope Analysis

M0, M1, …, Mn refers to the isotopologues containing n heavy atoms in a molecule. The stable isotope distribution of individual metabolites was measured by GC-MS as described above. The isotopologue enrichment or labeling in this work refers to the corrected isotope distribution ^24,25^.

### Gene expression of *Gls* from chondrosarcoma patient samples

We used retrospective data from RESOS INCA network of bone, collected from 102 cartilage tumors in different French hospitals. Multiple samples may be taken from each tumor and sequenced. More details about the experiment, such as RNA isolation method and profiling, can be found the original paper [40]. For our purposes, we utilized IDH mutation information to categorize samples into two groups: *IDH1* or *IDH*2 mutation (n=46) and *IDH1* and *IDH2* wild type (n=98). We examined several gene expression levels across these two groups and reported adjusted p-value for GLS result (with Benjamini–Hochberg correction). All the data processing and calculation was performed using Bioconductor docker (devel), with R version 4.0.3 and Bioconductor version 3.13.

### Quantification and Statistical Analysis

Statistical analyses were performed using Graphpad Prism 9 software. Data were presented as mean±SEM, mean±SD, or mean±95% as specified in each figure.

Statistical significance was determined by two tailed student-t test or one-way or two-way ANOVA with multiple comparisons test as specified in each figure.

## Acknowledgement

Research reported in this publication was supported by the National Institute of Arthritis and Musculoskeletal and Skin Diseases of the National Institutes of Health under award R01AR066765. The content is solely the responsibility of the authors and does not necessarily represent the official views of the National Institutes of Health.

## Conflict of Interests

The authors declare no conflicts of interests.

---

**Supplementary Fig 1:**
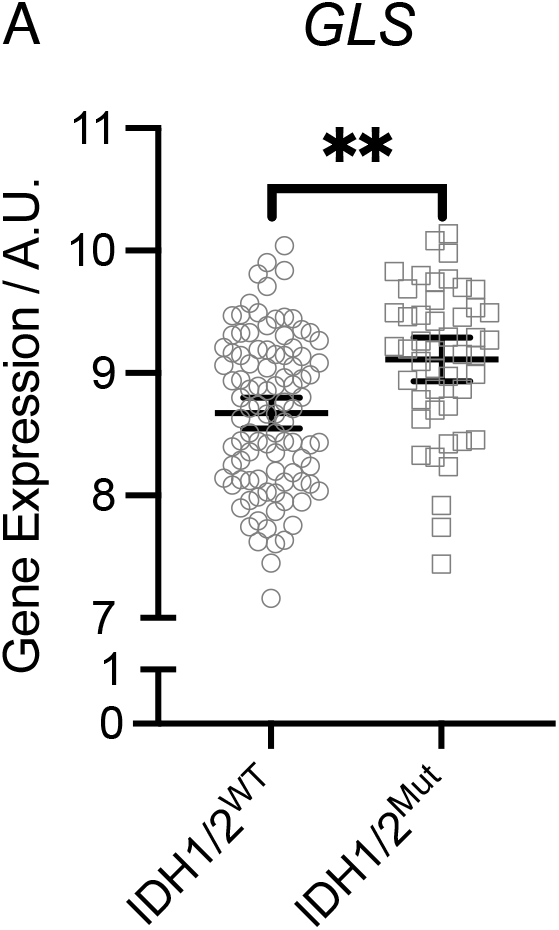
*GLS* was upregulated in chondrosarcoma tumors with *IDH1* or *2* mutations. Relative gene expression of *GLS* in human chondrosarcoma patient samples with wildtype or mutant *IDH1* or *IDH2.* mean±95% CI are shown. **padj<0.05.

**Supplementary Fig 2:**
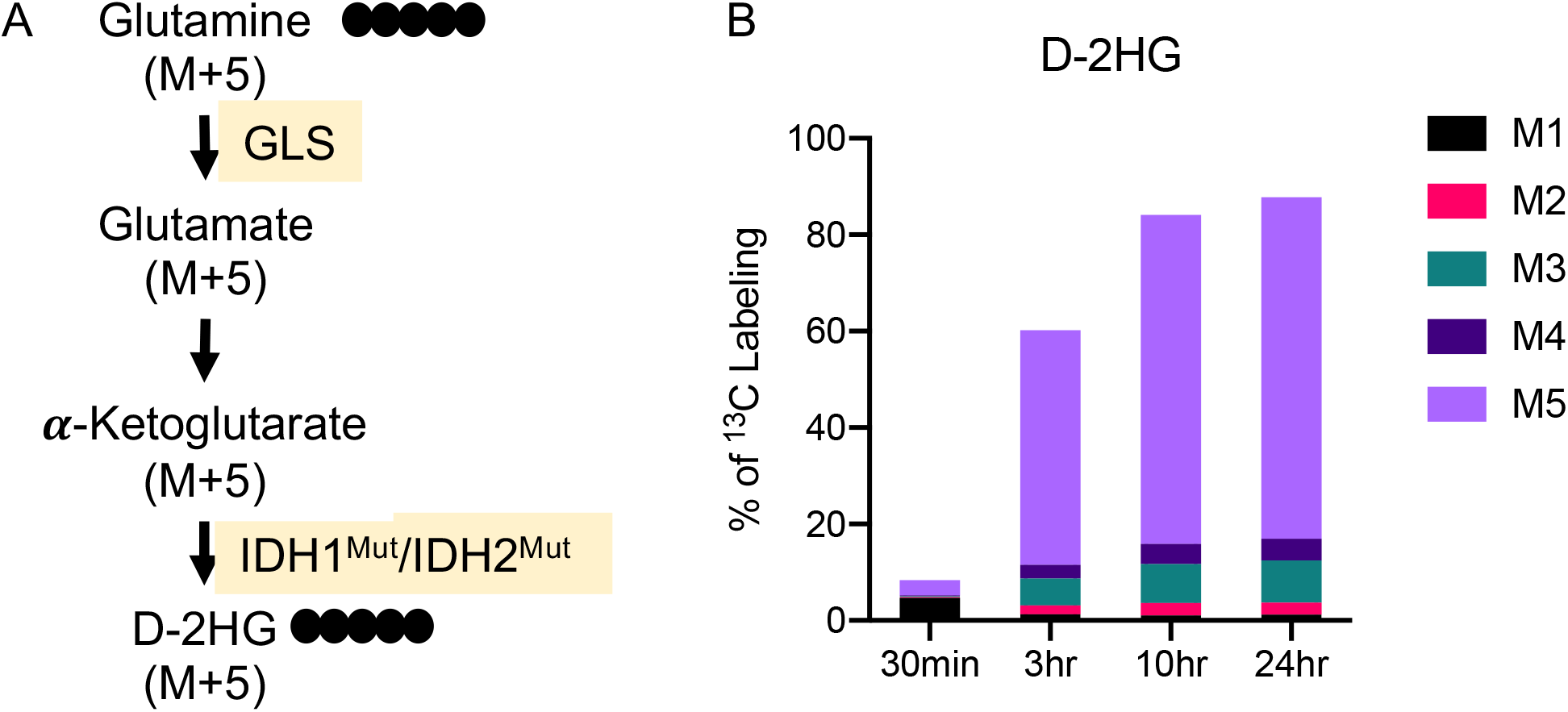
Glutamine was the primary source for D-2HG production in chondrocytes with *Idh1* mutation. (A) Graphical depiction of tracing glutamine metabolism using ^5^C5-glutamine. Filled circles indicate ^13^C and open circles indicate ^12^C. (B) AdCre;*Idh1^LSL/+^* chondrocytes used glutamine for D-2HG production. Percentage of ^13^C labeling in D-2HG at different time points.

**Supplementary Fig 3:**
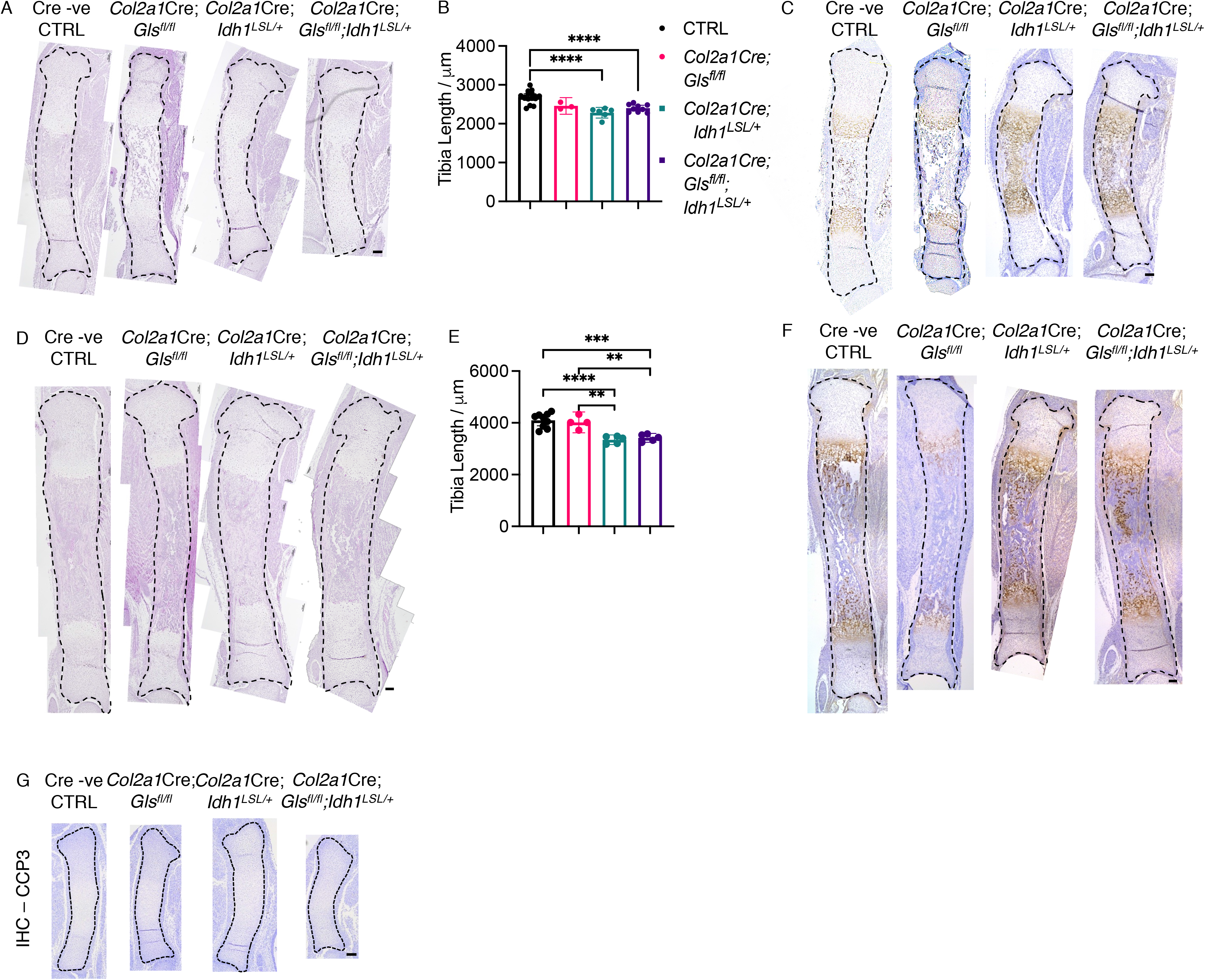
*Col2a1*Cre*;Idh1^LSL/+^* and *Col2a1*Cre;*Gls^fl/fl^;Idh1^LSL/+^* animals displayed defects in bone growth and hypertrophic chondrocyte differentiation. (A) H&E staining of murine tibias at E16.5. (B) Quantification of tibia length based on H&E staining at E16.5. (C) Immunohistochemistry of Col X of murine tibias at E16.5. (D) H&E staining of murine tibias at E18.5. (E) Quantification of tibia length based on H&E staining at E18.5. (F) Immunohistochemistry of Col X of murine tibias at E18.5. (G) Immunohistochemistry of cleaved caspase 3 of murine tibias at E14.5. mean±SEM are shown. **p<0.01, ***p<0.001, ****p<0.0001 (ANOVA)

**Supplementary Fig 4:**
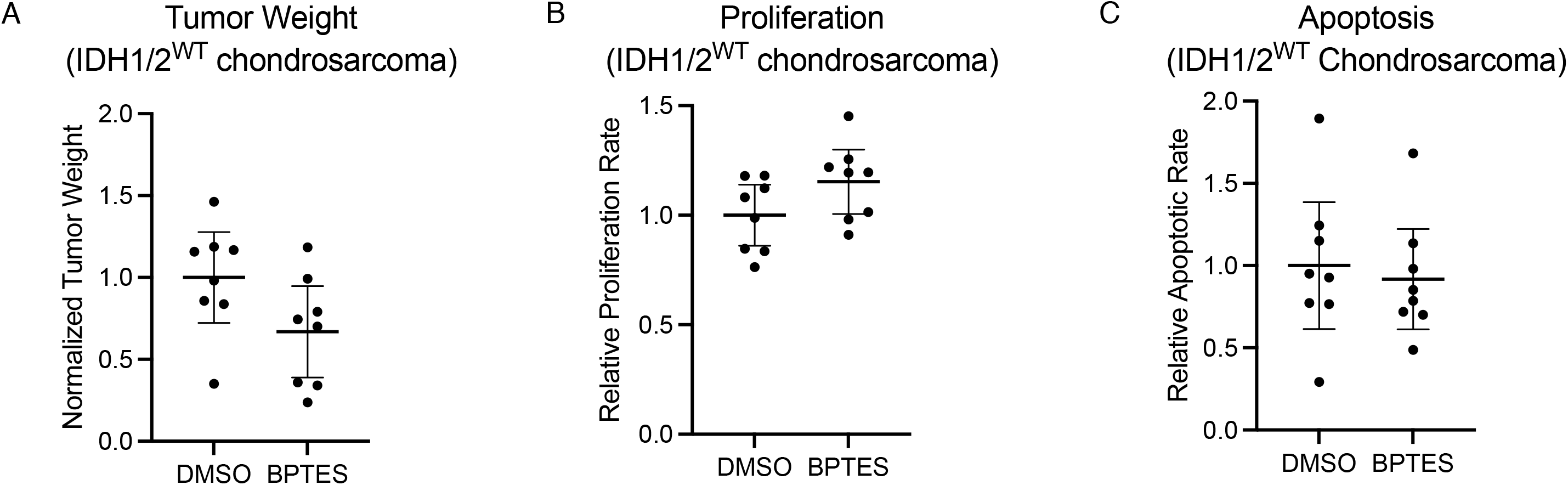
BPTES treatment did not cause significant change in tumor weight and cell viability of *IDH1* and *IDH2-*wildtype chondrosarcoma. (A) Relative tumor weight of xenografted *IDH1* and *IDH2-*wildtype chondrosarcoma tumors at the time of harvest. (B) Relative proliferation rate of each *IDH1* and *IDH2-*wildtype tumor determined by percentage of KI67 positive cells. (C) Relative apoptotic rate of each IDH1/2^WT^ tumor determined by percentage of cleaved caspase 3 positive cells. mean±95% CI are shown. Significance was determined by unpaired student t-test.

**Supplementary Fig 5:**
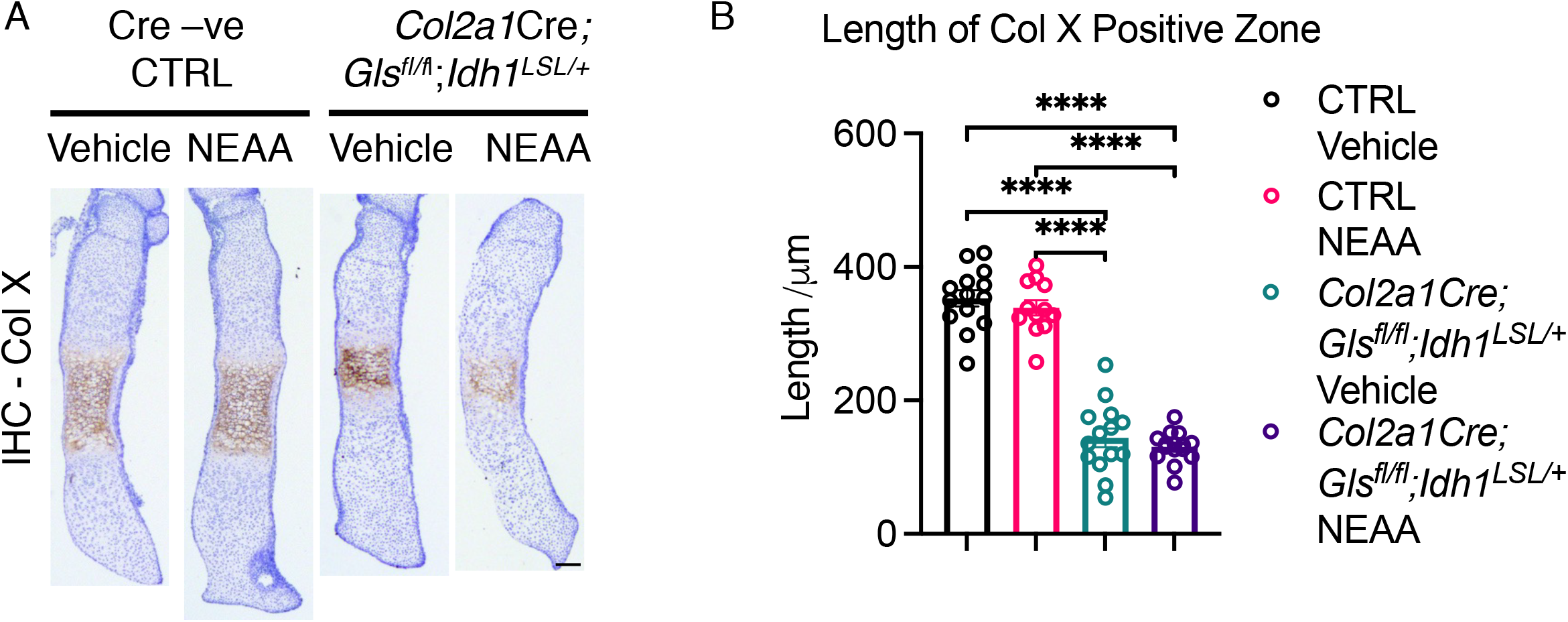
Supplementing non-essential amino acids did not recue defects in hypertrophic chondrocyte differentiation in *Col2a1*Cre;*Gls^fl/fl^;Idh1^LSL/+^* metatarsal organ culture. (A) Immunohistochemistry of Col X of metatarsal organ culture at E16.5. (B) Quantification of the length of Col X positive zone in metatarsal organ culture. mean±SEM are shown. ****p<0.0001 (ANOVA)

## Notes

https://www.ebi.ac.uk/arrayexpress/experiments/E-MTAB-7264/

https://www.ncbi.nlm.nih.gov/geo/query/acc.cgi?acc=GSE123130

### Competing Interest Statement

The authors have declared no competing interest.

